# Evaluation of endogenous and therapeutic 25-hydroxycholesterols in murine models of pulmonary SARS-CoV-2 infection

**DOI:** 10.1101/2022.09.12.507671

**Authors:** Michael B. Fessler, Jennifer Madenspacher, Paul J. Baker, Kerry L. Hilligan, Ehydel Castro, Julie Meacham, Shih-Heng Chen, Reed F. Johnson, Negin P. Martin, C.J. Tucker, Debabrata Mahapatra, Mark Cesta, Katrin D. Mayer-Barber

**Author notes:** **Correspondence to:** <u>Michael B. Fessler, M.D.</u>, National Institute of Environmental Health Sciences, 111 T.W. Alexander Drive, P.O. Box 12233, MD D2-01, Research Triangle Park, NC 27709, and <u>Katrin D. Mayer-Barber, PhD</u>, National Institute of Allergy and Infectious Diseases, Bldg 33, Room 2W10.A3, 33 North Drive, MSC 3206, Bethesda, MD, 20892, USA. Equal contribution (Michael B. Fessler, Katrin D. Mayer-Barber).

## Abstract

Oxysterols (i.e., oxidized cholesterol species) have complex roles in biology. 25-hydroxycholesterol (25HC), a product of activity of cholesterol-25-hydroxylase (CH25H) upon cholesterol, has recently been shown to be broadly antiviral, suggesting therapeutic potential against SARS-CoV-2. However, 25HC can also amplify inflammation and tissue injury and be converted by CYP7B1 to 7α,25HC, a lipid with chemoattractant activity via the G protein-coupled receptor, EBI2/GPR183. Here, using *in vitro* studies and two different murine models of SARS-CoV-2 infection, we investigate the effects of these two oxysterols on SARS-CoV-2 pneumonia. We show that while 25HC and enantiomeric-25HC are antiviral *in vitro* against human endemic coronavirus-229E, they did not inhibit SARS-CoV-2; nor did supplemental 25HC reduce pulmonary SARS-CoV-2 titers in the K18-human ACE2 mouse model *in vivo*. 25HC treatment also did not alter immune cell influx into the airway, airspace cytokines, lung pathology, weight loss, symptoms, or survival but was associated with increased airspace albumin, an indicator of microvascular injury, and increased plasma pro-inflammatory cytokines. Conversely, mice treated with the EBI2/GPR183 inhibitor NIBR189 displayed a modest increase in lung viral load only at late time points, but no change in weight loss. Consistent with these findings, although *Ch25h* was upregulated in the lungs of SARS-CoV-2-infected WT mice, lung viral titers and weight loss in *Ch25h*^−/–^ and *Gpr183*^−/–^ mice infected with the beta variant were similar to control animals. Taken together, endogenous 25-hydroxycholesterols do not significantly regulate early SARS-CoV-2 replication or pathogenesis and supplemental 25HC may have pro-injury rather than therapeutic effects in SARS-CoV-2 pneumonia.

## Introduction

In 2013, two groups identified 25-hydroxycholesterol (25HC), an oxysterol product of the enzyme cholesterol-25-hydroxylase (CH25H), as broadly antiviral (1, 2). All enveloped viruses tested to date, among them, influenza A, herpes simplex virus-1, varicella zoster virus, murine gamma herpes virus, HIV, vesicular stomatitis virus, hepatitis viruses B and C, and Ebola virus, are inhibited by 25HC, with IC_50_ in the nanomolar to low-micromolar range (3). 25HC blocks viral fusion by modifying host cell membranes and potentially by inhibiting the cholesterol-synthetic transcription factor SREBP2 (3). Additional posited mechanisms include effects on virus capsid disassembly, genome replication, protein expression, and cellular egress (2).

CH25H is expressed by airway epithelium (4) and alveolar macrophages (5, 6) in mice and humans and induces robust extracellular release of 25HC after upregulation by virus and interferons (1). Systemic treatment with exogenous 25HC to augment this native response effectively treats viral infections, including pneumonia, in mice, pigs, and non-human primates (2, 7, 8). Moreover, exogenous 25HC is anti-inflammatory through inhibiting inflammasomes (9) and accelerates resolution of lung inflammation in mice through activation of Liver X Receptor (LXR) (5). Reports such as these have collectively suggested potential for 25HC as a therapeutic for viral pneumonia. However, other reports that 25HC amplifies pro-inflammatory signaling and that *Ch25h*^−/–^ mice have attenuated lung pathology during influenza A pneumonia (10) have suggested possible untoward effects of 25HC *in vivo*. Further complicating questions of mechanism, 25HC is converted by the enzyme CYP7B1 into 7α,25-dihydroxycholesterol (7α,25HC), a ligand for Epstein Barr Virus-induced gene 2 (EBI2, encoded by *Gpr183*), a G protein-coupled receptor that can promote migration of several immune cell types to the lung (4, 6, 11, 12).

Of interest, it was recently reported that 25HC inhibits SARS-CoV-2 infection in cell lines (13-17). One group also reported that treatment of mice with 25HC reduced lung viral load 3 days post-infection (p.i.) with mouse-adapted SARS-CoV-2, but no further outcomes were presented (16), leaving many questions unaddressed. Here, we show that while both 25HC and enantiomeric (ent)-25HC inhibit cellular infection by endemic human coronavirus (hCoV)-229E they failed to limit SARS-CoV-2 infection. Moreover, during *in vivo* SARS-CoV-2 infections, *Ch25h*-deficient mice were able to control viral replication and transgenic mice expressing human ACE2 under the epithelial K18 promotor (K18-ACE2 mice) treated with supplemental 25HC did not exhibit changes in viral titers. 25HC treatment had no effect on airway immune response, lung histopathology, morbidity, or mortality but did increase plasma chemokines as well as bronchoalveolar lavage fluid (BALF) albumin, a metric of microvascular injury (18). *Gpr183-*deficient mice displayed no significant changes in lung viral loads or weight loss, thus arguing against a role for the downstream lipid, 7α,25HC. Taken together, our findings suggest that 25HC is not therapeutic during SARS-CoV-2 pneumonia and that GPR183 and CH25H are dispensable for early viral control *in vivo*.

## Materials and Methods

### Reagents

25HC was from Sigma Aldrich (St. Louis, MO) and ent-25HC was custom-synthesized by Avanti Polar Lipids (Birmingham, AL). The GPR183-specific antagonist NIBR189 was from Tocris (Bio-Techne Corp, Minneapolis, MN).

### Mice

Male and female B6.Cg-Tg(K18-ACE2)2Prlmn/J (#034860, Jackson Laboratory), C57BL/6 (B6; Jackson Laboratory), *Ch25h*^−/–^ (#016263, Jackson Laboratory) (5), and *Gpr183*^−/–^ ((19), kindly provided by Vanja Lazarevic) mice, ∼22-26g in weight, were used. The light/dark cycle was set at 12/12 hours, and mice were fed Purina Lab Diet #5002 and provided water *ad libitum*. Animal care and housing met AAALAC International guidelines, the Guide for Care and Use of Laboratory Animals (National Research Council), and requirements as stated by the U.S. Department of Agriculture through the Animal Welfare Act. All experiments were performed in compliance with an animal study proposal approved by the NIAID Animal Care and Use Committee or the NIEHS Animal Care and Use Committee.

### *In vivo* murine SARS-COV2 viral infections and treatments

In some studies, K18-hACE2 mice were treated i.p. with 50 mg/kg 25HC or vehicle (hydroxypropylbeta-cyclodextrin [Cyclo Therapeutics, Inc.]) at -4h in relation to viral infection, and every 24h thereafter until sacrifice. In other studies, K18-hACE2 mice were i.p. injected with 0.1mg/kg or 0.5 mg/kg NIBR189 on days -1, +1, and +3 p.i. K18-hACE2 mice were infected i.n. with 35 μL of neat USA-WA1/2020 SARS-CoV-2 (BEI), with target dose of 1+E03 TCID_50_/mouse. SARS-CoV-2 USA-WA-1 (WA1/2020), GenBank MN985325.1 was propagated on VEROE6 TMPRSS2 cells (20). The virus sequence was confirmed by Illumina sequencing and did not differ from the parental virus, except for S6L in E, T7I in M, and S194T in N (virus used in 25HC treatment experiments) and for the following SNPs compared to reference sequence: C23525T, C26261T, C26542T, and T28853A (virus used in NIBR189 experiments). All experiments with SARS-CoV-2 were performed at BSL-3. B6, *Ch25h*^−/–^, and *Gpr183*^−/–^ mice were infected i.n. with with 3.5×10^4^ TCID_50_ SARS-CoV-2 hCoV-19/South Africa/KRISP-K005325/2020 beta variant of concern (Pango lineage B.1.351, GISAID reference: EPI_ISL_860618, BEI resources) in 35μl. Exposure doses were confirmed by TCID_50_ (median tissue culture infectious dose) assay of remaining stock in VeroE6 cells (CRL-1586; American Type Culture Collection). Prior to inoculation, mice were anesthetized with ketamine (80-100 mg/kg) and xylazine (5-10 mg/kg) or isoflurane.

### Symptom scoring

Mice were weighed and scored daily for symptoms. A value of 1 was assigned for any of the following signs and individual values summed to yield a composite score: rough coat, hunched posture, lethargy, prostrate status, labored breathing, decreased breathing, respiratory distress, increased breathing, eye discharge.

### Animal harvests

In some studies, at time of sacrifice, mice underwent plasma collection and bronchoalveolar lavage (BAL) with 1x PBS (5). The cell count was quantified by Guava® easyCyte™ flow cytometer and differentials determined manually after Wright Giemsa staining of cytospin. In survival studies, euthanasia was performed for: (i) respiratory distress; (ii) prostration; (iii) unresponsive status to touch or external stimuli; (iv) ≥30% body weight loss; or (v) ≥25% body weight loss plus lethargy, decreased activity, or ataxia.

### Cytokine and albumin analysis

Cell-free BAL fluid (500x g, 5 min) or plasma was analyzed using ELISA kits for CCL2, CXCL1, CXCL10, IL-6, CCL5, and TNFα (Thermo Fisher Scientific) and albumin (Abcam), per manufacturer’s instructions.

### RNA isolation and qPCR

RNA was isolated by RNeasy 96 QIAcube HT kit with a DNAse step. Complementary DNAs (cDNA) were generated using standard methods. Real-time PCR was performed in triplicate with Taqman PCR Mix (Applied Biosystems) in the HT7900 ABI sequence Detection System (Applied Biosystems). Predesigned primers were purchased from Applied Biosystems (*Ch25h* (Mm00515486_s1), *Gapdh* (Mm99999915_g1) as a housekeeping gene for normalization). For quantitation of viral load, the N1 nucleocapsid gene was quantified, using serial dilutions of reference standard input RNA to achieve a linear relationship between threshold cycle and target RNA quantity. Forward primer: 5’-GAC CCC AAA ATC AGC GAA AT-3’. Reverse primer: 5’-TCT GGT TAC TGC CAG TTG AAT CTG-3’. Probe: 5’-FAM-ACC CCG CAT TAC GTT TGG TGG ACC-NFQ-MGB-3’ (Thermo Fisher Scientific).

### TCID_50_ assay from mouse lung

Lungs were harvested into tubes containing 600μL-1mL sterile PBS and 1mm silicon beads before being homogenized using a Precelleys Tissue Homogenizer (Bertin Instruments). 10-fold serial dilutions of lung homogenate in DMEM containing 2% FCS were plated in triplicate or quadruplicate on VeroE6 cells (CRL-1586; American Type Culture Collection) in 96-well cluster plates seeded with 2.5×10^4^ cells per well the previous day. Plates were stained with Crystal Violet solution (Sigma) after 96-168h to assess cytopathic effect (CPE). Viral titers were determined using the Reed-Muench or Spearman Kärber method.

### Histopathologic analysis

Tissues were fixed in 10% neutral-buffered formalin for a minimum of 48h and transferred into 70% ethanol for a minimum of 72h to inactivate virus, processed for paraffin, embedded, sectioned (5 μm), and stained with H&E. Slides were scanned using an Aperio ScanScope XT Scanner (Aperio Technologies, Inc., Vista, CA). Lung histopathology was scored in a blinded fashion by a board-certified veterinary pathologist, using an equally weighted composite score for: cellular inflammation, septal thickening, type II alveolar epithelial cell hyperplasia, airway epithelial hyperplasia/hypertrophy, necrosis, and hemorrhage (all scored 0 [none] – 4 [severe]).

### hCoV-229E plaque reduction assay

MRC-5 monolayers in 6-well plates (1×10^6^ cells/well) were pretreated overnight with medium containing vehicle, 2 μM 25HC, or 2 μM ent-25HC. Monolayers were then inoculated for 1h with 100 μL of medium containing ∼100 PFU of hCoV-229E. The infected monolayers were left for 6-8 days in a 35°C incubator under 0.8% methylcellulose (Millipore Sigma) overlay (in EMEM medium with 10% FBS, 10 mM HEPES, 2 mM L-Glutamine, 100 units penicillin, 0.1 mg streptomycin) containing vehicle control, 2 μM 25HC or 2 μM of ent-25HC, respectively, until plaques became visible. The cell monolayers were then stained with 2% crystal violet in 10% formalin in PBS for at least 2h. The stained monolayers were washed and air-dried, and clear plaques were counted manually under the light microscope.

### SARS-CoV-2 plaque reduction assay

VeroE6 expressing TMPRSS2 cells (20) in DMEM 10% FBS were plated at 400,000 cells/well in 12-well dishes and incubated overnight at 37°C, 5% CO_2_. Compounds in DMSO were diluted in DMEM with 2% FBS with a fixed concentration of 0.05% DMSO. Remdesivir (Adipogen Life Sciences, San Diego, CA) was used as a positive control. Two-fold dilutions of antibody were made and tested. Cells were washed twice in DMEM+2%FBS, then pre-treated with compounds for 2h prior to infection at no DMSO, 0.05% DMSO, 2.5, 5, or 10μM of compound at 37°C, 5% CO_2_. Forty plaque forming units of virus in 200uL of DMEM+2% FBS was mixed with 100uL of media with no DMSO, 0.05% DMSO, 2.5, 5, or 10μM of compound. The virus+ compound mixtures were briefly vortexed and incubated at 37°C for 30 minutes. Cells were washed twice with DMEM+2% FBS, the second wash was removed and 100uL of the virus+ compound mixture was added to duplicate wells, incubated at 37°C, 5% CO_2_ for 1hr. Cells were overlayed with 0.6% methylcellulose and incubated for 72h at 37°C, 5% CO_2_. At 72h post-infection, the overlay was removed and the samples were fixed with crystal violet+ neutral buffered formalin (0.1% crystal violet, 20% ethanol v/v+10% NBF v/v) for 30min at room temperature. Plaques were enumerated and compared to no DMSO sample.

### SARS CoV-2 yield reduction assay

VeroE6 cells expressing TMPRSS2 (20) were plated in DMEM 10% FBS at 200,000 cells/well in 12-well dishes and incubated overnight at 37°C, 5% CO_2_. Cells were washed twice in DMEM+2%FBS, pre-treated with compounds for 24h prior to infection at no DMSO, 0.05% DMSO, 2.5, 5, or 10μM of compound at 37°C, 5% CO_2_ in DMEM+2% FBS. Cells were washed twice in DMEM+2%FBS, infected at an MOI=0.1 for 1hr at 37C, 5% CO_2_. Cells were then washed twice in DMEM+2%FBS, then media with no DMSO, 0.05% DMSO, 2.5, 5, or 10μM of compound at 37°C, 5% CO_2_ was added to the wells. Media was collected at 24h post-infection and assayed by plaque assay. For plaque assays, 400,000 cells per well were plated in 12-well dishes and incubated overnight at 37°C, 5% CO_2_ and washed twice in DMEM+2%FBS prior to the plaque assay. Samples were serially 10-fold diluted in DMEM+2% FBS and added to the cells followed by incubation for 1 hr. At the end of were washed twice in DMEM+2%FBS. At the end of 1h, cells were overlayed with 0.6% methylcellulose and incubated for 72h at 37°C, 5% CO_2_. At 72h post-infection, the overlay was removed and the samples were fixed with crystal violet+ neutral buffered formalin (0.1% crystal violet, 20% ethanol v/v+10% NBF v/v) for 30min at room temperature. Plaques were enumerated and compared to no DMSO sample.

### Viral plaque image analysis

For analysis of plaque size, brightfield images were captured on a Zeiss AxioObserver Z1 microscope (Carl Zeiss Inc, Oberkochen, Germany) using a Fluar 5X objective with a halogen light source. Transmitted light was collected with a Zeiss AxioCam IC color camera with an exposure of 2.04ms. A full tiled image of each well of a 6 well multiwell plate was acquired and stitched using the Zeiss Zen Blue software. These stitched images were then imported into FIJI (ImageJ version 1.53c) where a macro was created to automate analysis. The custom macro extracted the green component of the RGB image, applied a Gaussian filter, set a threshold to identify the plaques, and then used Analyze Particles to count and measure the size of each plaque.

### Statistical analysis

Analysis was performed using GraphPad Prism software. Data are represented as mean ± SEM. Two-tailed Student’s *t* test was applied for comparisons of 2 groups and ANOVA for comparisons of more than 2 groups. Survival was evaluated by log-rank test. For all tests, *P*< 0.05 was considered significant.

## Results

### 25HC inhibits human endemic coronavirus infection but not SARS-CoV-2 *in vitro*

25HC is broadly antiviral against enveloped viruses, reportedly inhibiting virus fusion by altering cholesterol content of the host cell membrane (3). 25HC may sterically redistribute free cholesterol by incorporation into host membranes, although it has also been posited to reduce membrane free cholesterol through direct activation of the enzyme acyl-coenzyme A:cholesterol acyltransferase (17). Specific protein-binding interactions are typically not preserved in enantiomeric (ent) versions of lipids (1). We started with testing 25HC for efficacy against a common cold coronavirus. To distinguish the mode of mechanism, we pretreated MRC-5 cells with 25HC, ent-25HC, or vehicle and then infected them with the endemic human coronavirus (hCoV)-229E. As shown in **Figure 1A**, 25HC and ent-25HC both significantly reduced viral plaque number, with 25HC exhibiting a much more robust effect. 25HC and ent-25HC also reduced plaque size, suggestive of post-entry effects on replication (1), with 25HC again displaying a more marked effect (**Figure 1B**). Taken together, these findings suggest that 25HC antagonizes hCoV-229E through both protein binding-dependent and -independent mechanisms.

**Figure 1.**
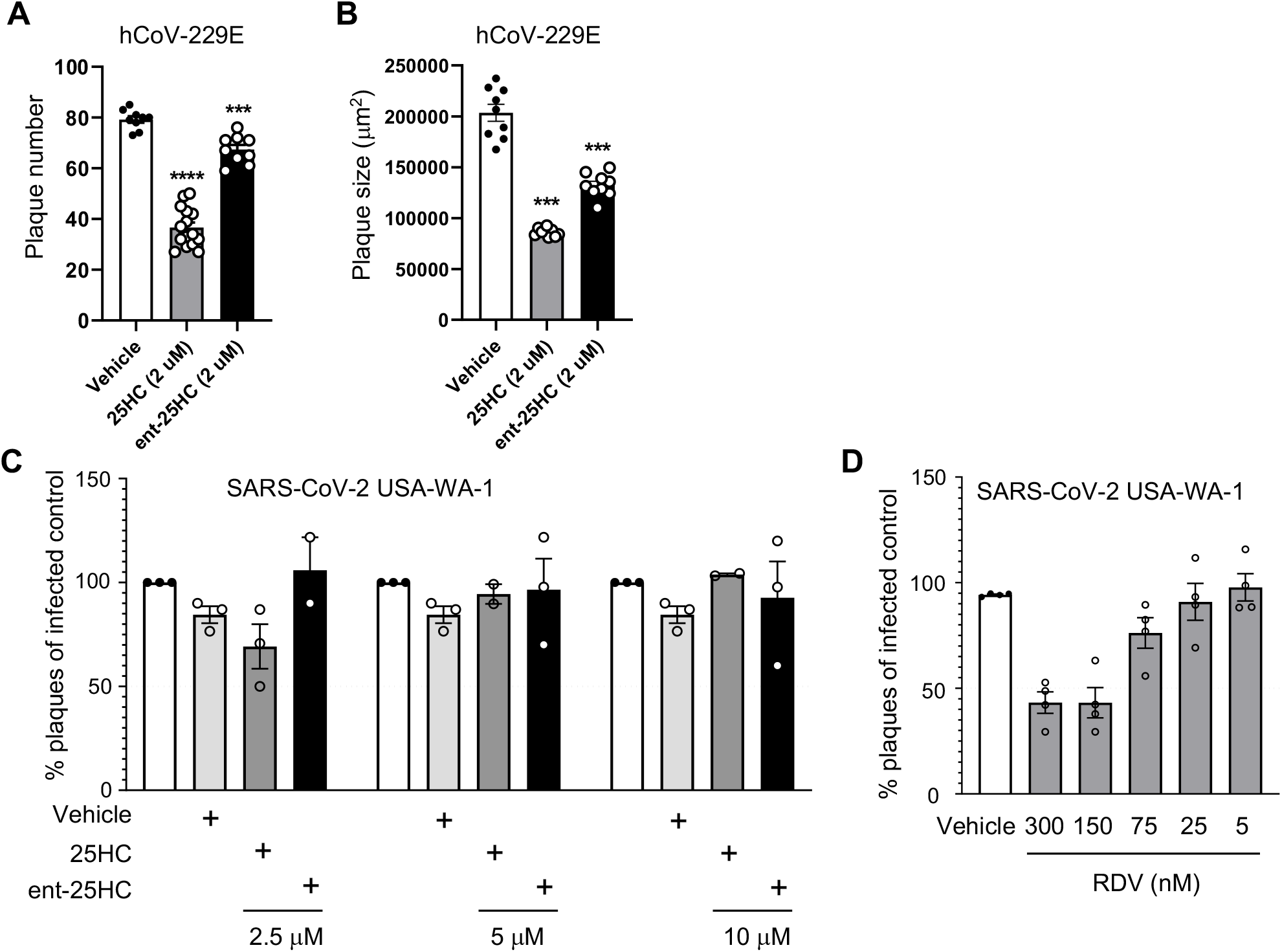
25HC antagonizes coronavirus infection *in vitro*. (**A-B**) MRC-5 cells were treated with 2 μM 25HC, 2 μM enantiomeric (ent)-25HC, or vehicle overnight, infected with hCoV-229E, and then evaluated for plaque number (A) and plaque size (B). (**C**) TMPRSS2-expressing VeroE6 cells were treated overnight with the indicated concentrations of oxysterols or vehicle, infected with SARS-CoV-2, and then evaluated for plaque number. (**D**) As a positive control to confirm assay performance, remdesivir (RDV) at a range of concentrations was tested for viral plaque reduction under the same conditions as in panel *C*. Data are mean +/- s.e.m. and are representative of 2-3 independent experiments. ***, P<0.001; ****, P<0.0001

We next tested 25HC and ent-25HC for antagonism against SARS-CoV-2 in VeroE6 cells stably transfected with TMPRSS2. Contrary to the case with hCoV-229E, neither 25HC nor ent-25HC exhibited a significant effect on SARS-CoV-2 replication at concentrations up to 10 μM, as indicated by both plaque reduction assay and yield reduction assay (**Figures 1C-D, E1**).

### Neither native CH25H nor exogenous 25HC alters viral replication in lungs of SARS-CoV-2-infected mice

We next tested the role of CH25H and 25HC in pulmonary host defense against SARS-CoV2 *in vivo*. We found that *Ch25h* was robustly upregulated in the lungs of K18-hACE2 transgenic mice (21) at 48h and 72h p.i. with SARS-CoV-2 (**Figure 2A**). Despite this, *Ch25h*^−/–^ mice infected with the beta variant of SARS-CoV-2 (B.1.351) had equivalent viral load in the lungs and equivalent weight loss compared to WT mice 3 days p.i. (**Figure 2B-C**). To test whether supplemental 25HC might nonetheless have antiviral effect *in vivo* during SARS-CoV-2 pneumonia, we treated K18-hACE2 transgenic mice i.p. with 50 mg/kg/d 25HC (or vehicle), a 25HC dosing regimen that has been reported to be antiviral against HIV and Zika virus in mice (2, 7) and to accelerate resolution of lung inflammation in mice (5). However, exogenous 25HC, compared to vehicle (**Figure 3A**), did not significantly alter viral load in the lung, as assessed either by qPCR (**Figure 3B**) or plaque assay (**Figure 3C**), nor did it modify clinical symptom scores or weight loss (**Figure E2**).

**Figure 2.**
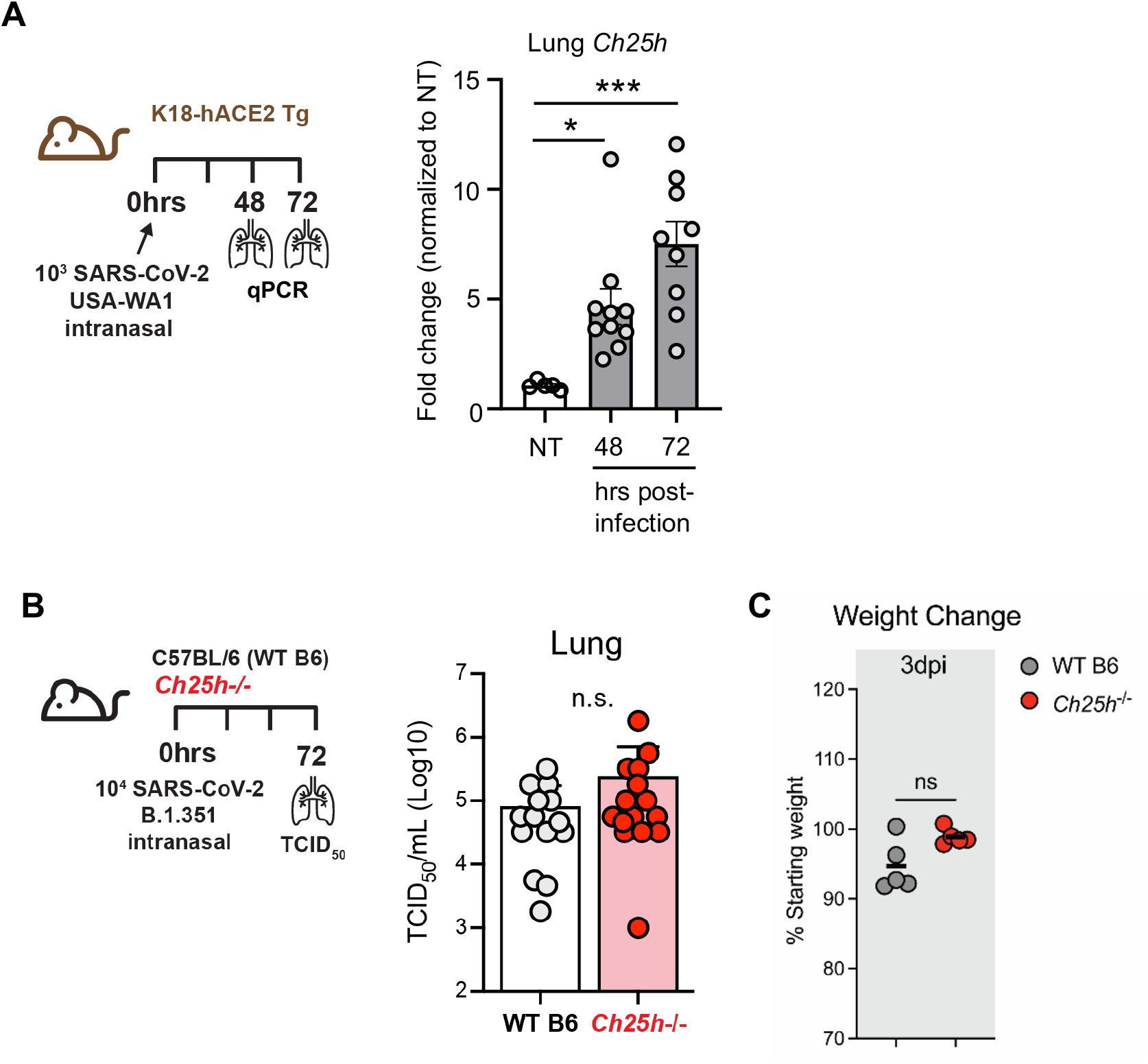
*Ch25h* is induced by SARS-CoV-2 in mouse lung but does not regulate viral clearance. (**A**) K18-hACE2 mice were left non-treated (NT) or infected with SARS-CoV-2 USA-WA1/2020. As per scheme on left, *Ch25h* expression was quantified in lung homogenates by qPCR at the indicated time points p.i. (N=5-10/condition). (**B**) As per scheme on left, WT or *Ch25h*^−/–^ mice were infected i.n. with SARS-CoV2 B.1.351 and lung viral load quantified by TCID_50_ assay on day 3 p.i. (N=15/genotype, 3 independent experiments). (**C**) Weight loss at day 3 p.i. was indexed to baseline weight (N=5/genotype). Data are mean +/- s.e.m. and are representative of 2-3 independent experiments. *, P<0.05; ***, P<0.001.

**Figure 3.**
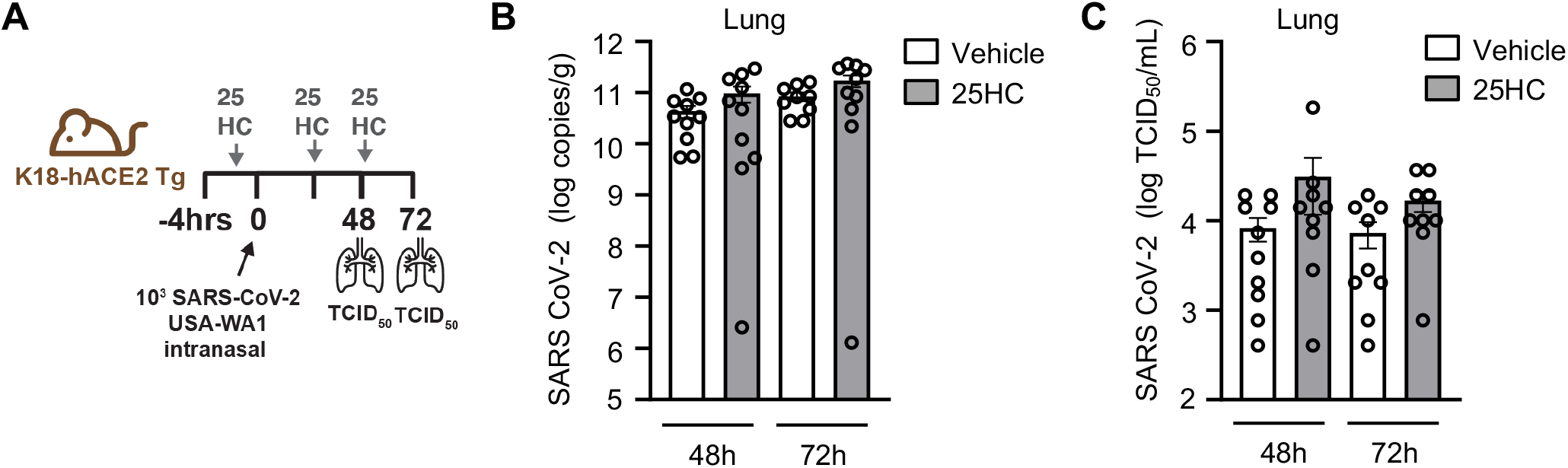
25HC treatment does not modify viral clearance during SARS-CoV-2 pneumonia. (**A**) K18-hACE2 mice were injected i.p. with 50 mg/kg/d 25HC (or vehicle) starting at -4h preceding i.n. inoculation with SARS-CoV-2, after which lungs were harvested at the indicated times p.i. and evaluated for viral load by qPCR (**B**) or plaque assay (**C**). N=9-10/condition. Data are mean +/- s.e.m.

### 25HC does not modify the pulmonary immune response to SARS-CoV-2

SARS-CoV-2, like other respiratory viruses, induces a time-dependent cellular immune response in the lungs (21). It is thought that acute lung injury, morbidity, and mortality during SARS-CoV-2 pneumonia stem largely from an overexuberant inflammatory response (21). Given this, we next profiled the pulmonary immune response of infected mice to investigate possible effects of 25HC. Total airway cell count in the BAL, as well as BALF concentrations of multiple pro-inflammatory cytokines and chemokines were all unchanged by 25HC treatment at 48-72h p.i. (**Figure 4A-B**). By contrast, several chemokines were elevated in the plasma of 25HC-treated mice 120h p.i. (**Figure 4C**), a time point by which significant morbidity has commenced in this model ((21) and Figure S2). Taken together, these findings indicate that treatment with 25HC neither alters viral clearance nor the immune/inflammatory response to SARS-CoV-2 in the mouse lung, but may be associated with elevated systemic levels of pro-inflammatory chemokines at later time points.

**Figure 4.**
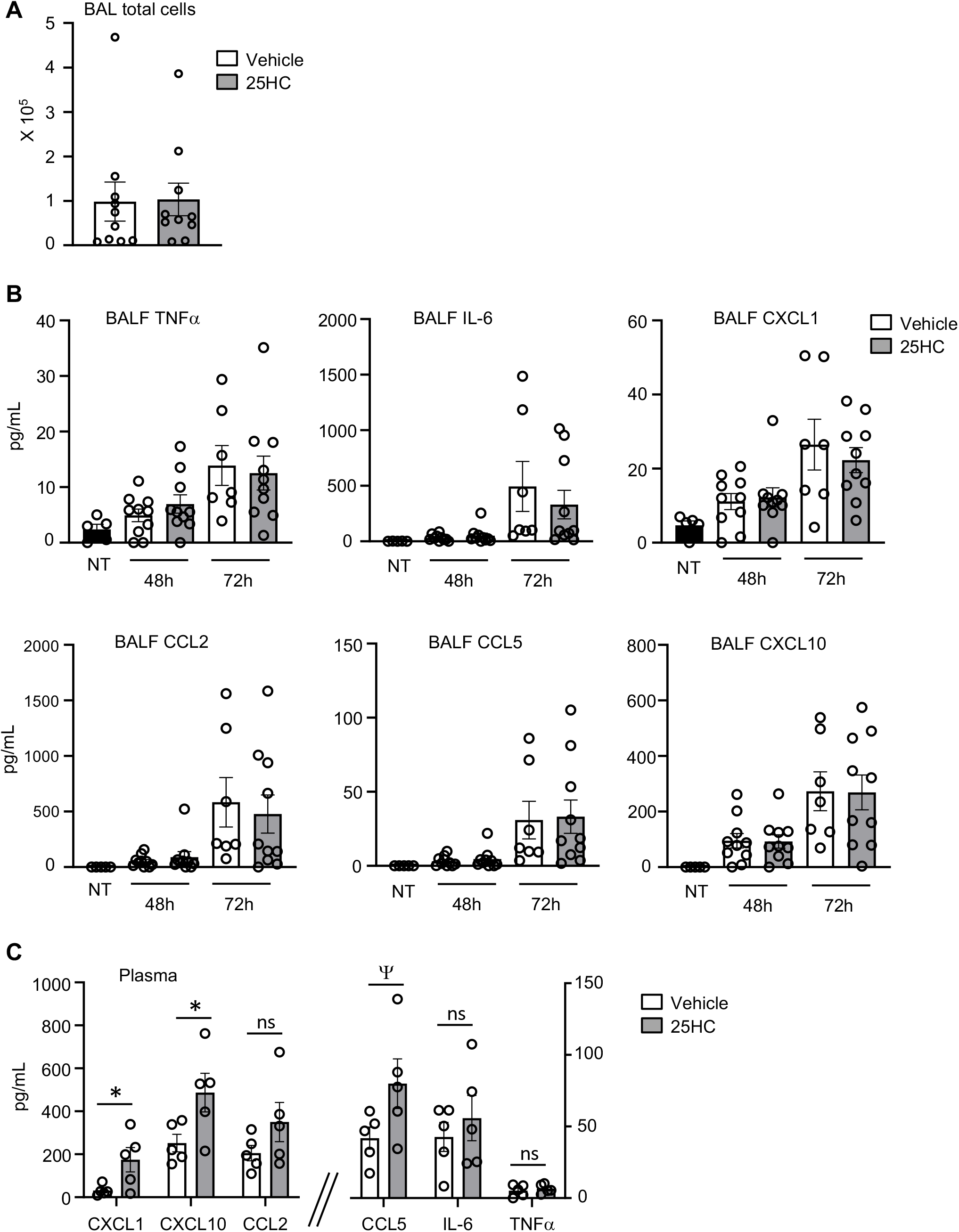
25HC treatment does not modify the inflammatory response during SARS-CoV-2 pneumonia. K18-hACE2 mice were either left non-treated (NT) or injected i.p. with 50 mg/kg/d 25HC (or vehicle) starting at -4h preceding i.n. inoculation with SARS-CoV-2. (**A**) Bronchoalveolar lavage (BAL) cells were quantified 48h post-infection (N=10/treatment). (**B**) Cytokines and chemokines were quantified in BAL fluid (BALF) at 48h and 72h post-infection (N=5-10/condition). (**C**) Plasma cytokines and chemokines were quantified at 120h post-infection (N=5/treatment). Data are mean +/- s.e.m. *, P<0.05; ^Ψ^, P=0.08. ns=nonsignificant.

### 25HC does not attenuate SARS-CoV-2-induced lung pathology and increases microvascular leak

SARS-CoV-2 induces severe acute lung injury that can lead to respiratory failure and death (21). Pulmonary vascular damage is a prominent feature of the pathogenesis and is thought to arise from direct viral infection or stimulation of endothelium (22). We found that, by 72-120h p.i., the lungs of K18-hACE2 transgenic infected mice exhibited severe neutrophilic and mononuclear cell infiltration that was associated with septal thickening, type II alveolar epithelial cell hyperplasia, airway epithelial cell hyperplasia, as well as focal necrosis and hemorrhage (**Figure 5A**). Comparable lung histopathology was observed in the lungs of vehicle- and 25HC-treated mice (**Figure 5A-B**). Of interest, 25HC-treated mice had elevated BALF concentrations of albumin (**Figure 5C**), an established metric of pulmonary microvascular injury (18). Regardless, 25HC had no effect on mortality (**Figure 5D**). Collectively, these findings suggest that supplemental 25HC does not overtly modify cellular histopathology in the lungs and has no effect on survival but may aggravate damage to the alveolocapillary barrier during SARS-CoV-2 pneumonia.

**Figure 5.**
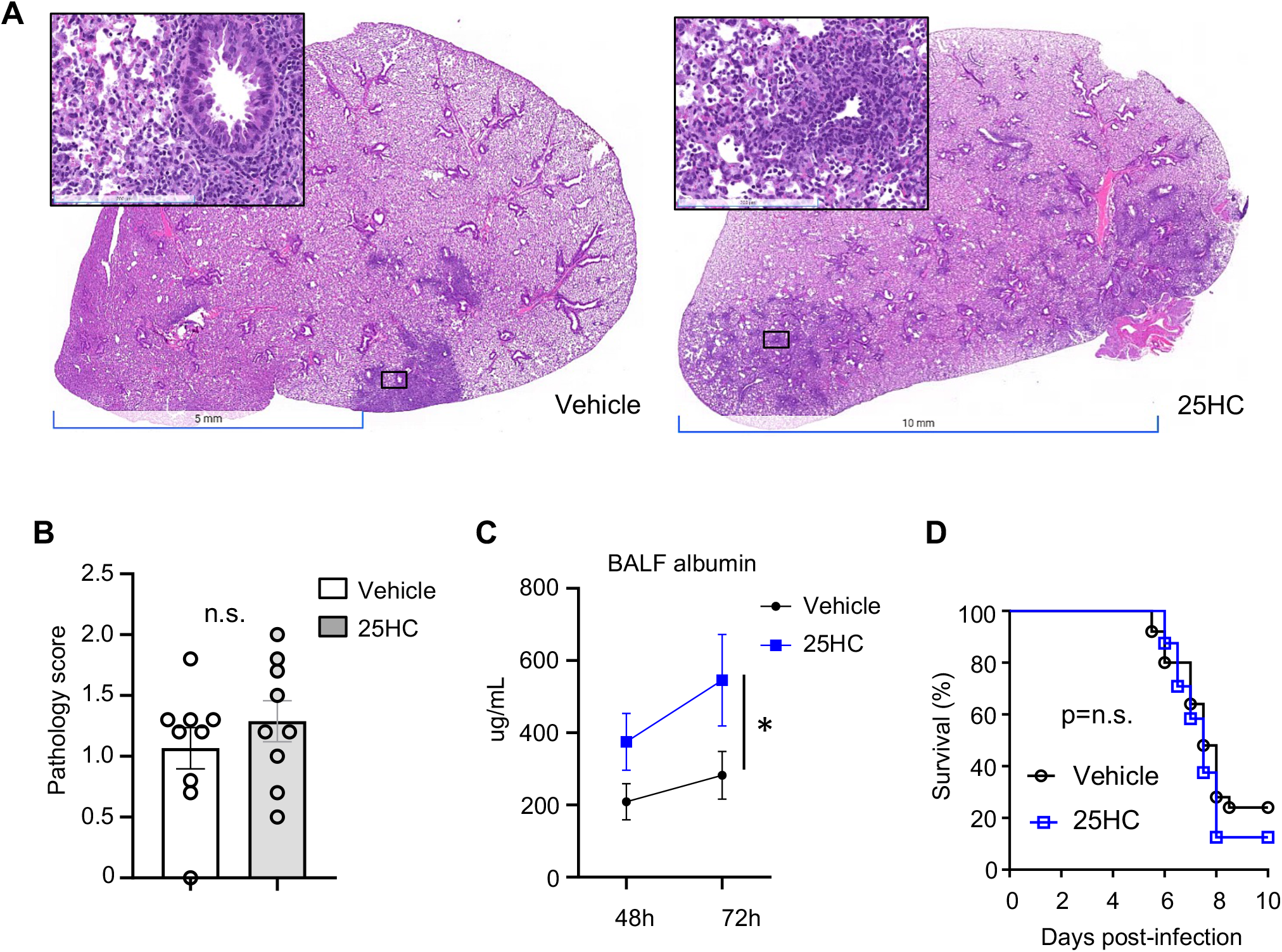
25HC treatment does not modify lung histopathology or mortality during SARS-CoV-2 pneumonia. K18-hACE2 mice were injected i.p. with 50 mg/kg/d 25HC (or vehicle) starting at -4h preceding i.n. inoculation with SARS-CoV-2. Lungs were harvested on day 5 p.i., fixed, stained with hematoxylin & eosin, and scored for histopathology. (**A**) Representative images and (**B**) composite pathology scores are shown (N=9/condition). (**C**) BALF albumin, an indicator of microvascular injury, was measured at the indicated times p.i. (N=5-10/condition). (**D**) Survival was monitored (N=25/treatment). Data are mean +/- s.e.m. and are representative of 2-3 independent experiments. n.s.= nonsignificant.

### Neither deletion nor pharmacological inhibition of *Gpr183*/EBI2 alter host defense after SARS-CoV-2 infection

After biosynthesis by CH25H, 25HC is converted by the enzyme CYP7B1 into 7α,25HC, an oxysterol ligand for Epstein-Barr virus-induced G protein-coupled receptor (GPR)2 (i.e., EBI2, encoded by *Gpr183)* (23). EBI2 is expressed by multiple leukocyte types and is thought to induce migration of Gpr183-expressing immune cells to the lung in response to 7α,25HC induced by allergens, cigarette smoke, and *M. tuberculosis* (4, 6, 11, 12). 7α,25HC levels are deficient in *Ch25h*^−/–^ mice (24). Given that *in vivo* studies of CH25H and 25HC may be confounded by downstream processing of 25HC into 7α,25HC (24, 25), we next investigated EBI2 in two models of SARS-CoV-2 pneumonia. First, *Gpr183*-null mice and WT controls were infected with SARS-CoV-2 B.1.351 (beta variant). Equivalent lung viral loads were found in both genotypes on day 3 p.i. (**Figure 6A**). Second, K18-hACE2 transgenic mice were treated with the EBI2 antagonist NIBR189 or vehicle and infected with SARS-CoV-2-WA1/2020. Treatment with 0.1 mg/kg NIBR189, a dose previously reported to be well tolerated for repeated treatments and pulmonary mode of action (26), did not change viral titers 3 days p.i. and was associated with only a modest increase in lung viral loads on day 5 p.i., but there were no accompanying changes in body weight **(Figure 6B**). Similar results were obtained using a 5-fold higher dose of inhibitor (not depicted). Taken together, these findings suggest that 7α,25HC or EBI2-mediated immune functions do not have a significant impact on host defense during SARS-CoV-2 pneumonia in mice.

**Figure 6.**
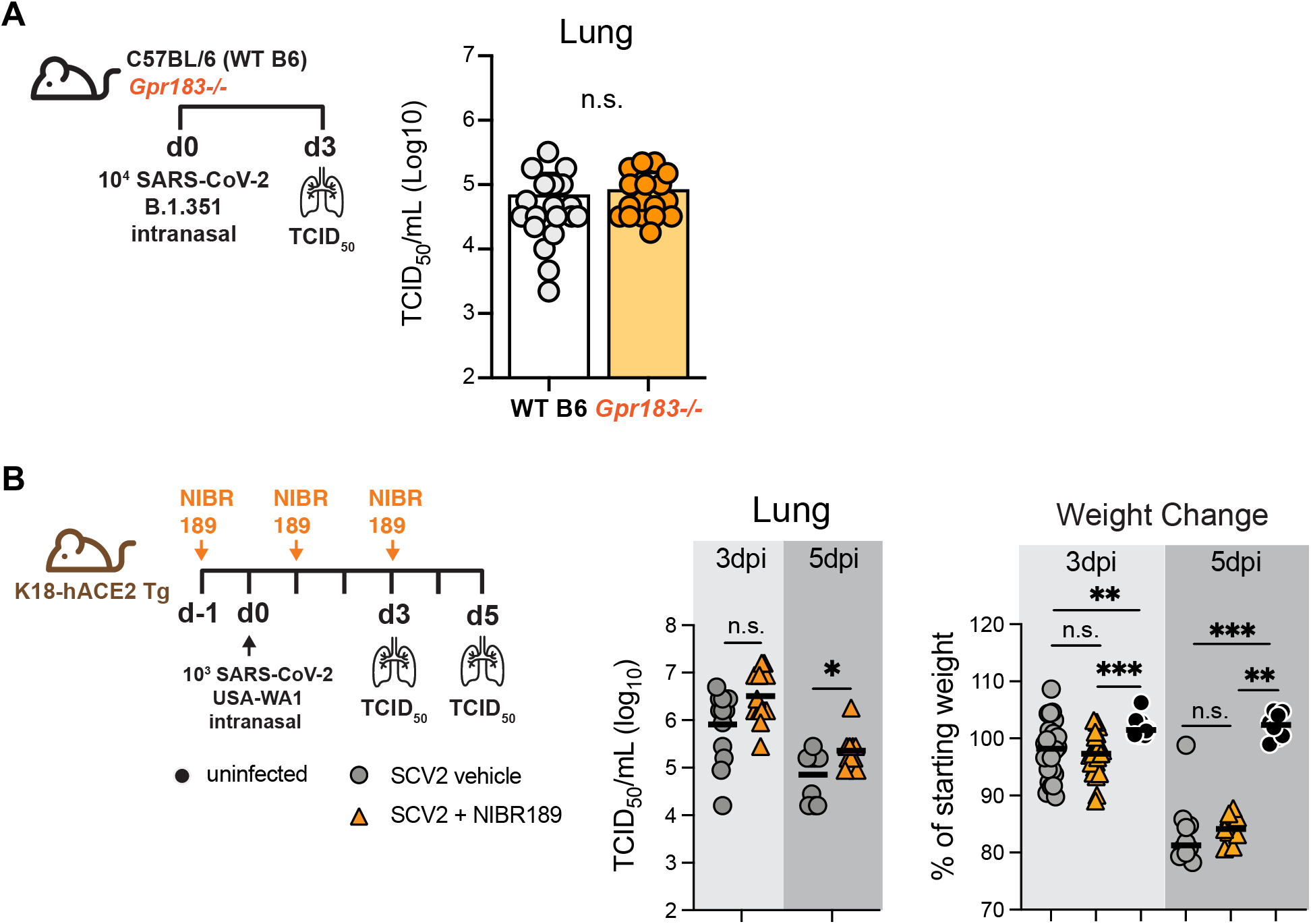
No effect of *Gpr183* deletion or EBI2 inhibition on SARS-CoV-2 clearance or morbidity. (**A**) As per scheme on left, *Gpr183*^−/–^ and WT (B6) mice were infected i.n. with a target dose of 3.5×10^4^ SARS-CoV-2 B.1.351. Viral load in lung was quantified on day 3 p.i. (N=18/genotype, 3 independent experiments (right). (**B**) As per scheme on left, K18-hACE2 transgenic mice were either left uninfected or treated i.p. with 0.1 mg/kg EBI2 (encoded by *Gpr183*) inhibitor NIBR189 (or vehicle control) and infected i.n. with a target dose of 1×10^3^ SARS-CoV2 USA-WA1/2020. Lung viral load (middle panel) and weight change (right panel) were quantified (N=7-20/condition, 2 independent experiments per time point). Data are mean +/- s.e.m. *, P<0.05; **, P<0.01; ***, P<0.001. n.s. = nonsignificant.

## Discussion

During the past several years, 25HC has been shown to inhibit infection by several viruses through multiple mechanisms and thus has been proposed as a therapeutic candidate (3, 8, 27, 28). 25HC reportedly blocks viral fusion by reducing free cholesterol in host cell membranes (15, 17), a mechanism that may either involve direct membrane intercalation of 25HC or activation by 25HC of the cholesterol-esterifying enzyme, acyl-coenzyme A (CoA):cholesterol acyltransferase (ACAT) (17). Additional antiviral mechanisms that have been proposed include inhibition of post-entry gene expression (1, 14), activation of the nuclear receptor LXR (29), inhibition of the replication organelle through interactions with oxysterol binding protein 1 (OSBP1) (30), and activation of the integrated stress response (31). While some of these mechanisms are proposed to involve direct binding interactions of 25HC with proteins (ACAT, OSBP1, LXR), other antiviral effects have been proposed to arise independently of 25HC-protein interactions (3). Our finding that 25HC had for more potent activity than ent-25HC against hCoV-229E suggests, consistent with a prior report (1), that 25HC is enantioselective, likely triggering protein-dependent effects at low concentrations which may be complemented by additional protein-independent mechanisms at higher concentrations.

Contrary to a few recent reports (14-17), we found no antiviral efficacy for 25HC against SARS-CoV-2 *in vitro*. This is despite our inclusion of 25HC concentrations higher than the EC50 for SARS-CoV-2 reported in these other publications (3.675 μM (16), 4.2 μM (14)), concentrations that are unlikely to be easily achievable *in vivo*. While we can only speculate on the reason for our different result, we presume that it reflects methodological differences, among them, choice of cell line (Calu-3, Caco-2 (17); HEK293-hACE2 (15)), virus strain (14, 15)), and timing of 25HC treatment (14)). Although not all reports have specified the vehicle used, we found no antiviral effect of 25HC for SARS-CoV-2 in either DMSO or ethanol (not depicted).

*In vivo*, we also did not detect any effect of supplemental 25HC or CH25H deletion on multiple disease measures during SARS-CoV-2 pneumonia. Recently, it was reported that 25HC, delivered intragastrically at 100 mg/kg/d starting 1 day prior to infection, reduced lung viral load of the mouse-adapted SARS-CoV-2 strain MASCp6 in Balb/C mice (16). Technical differences, including 25HC dose, 25HC route, virus strain, and mouse strain may possibly account for the divergence in our findings. Our dosing regimen (50 mg/kg/d i.p. in β-cyclodextrin vehicle) was previously reported to reduce HIV viral load in humanized mice (2) and to reduce Zika virus load and mortality in Balb/C mice (7). We have also previously shown that this 25HC regimen induces LXR target genes in mouse lung and is therapeutic against lung inflammation by enhancing efferocytic function of alveolar macrophages (5). This latter finding suggests that systemically dosed 25HC penetrates the airspace in biologically active form. Further supporting this, it was recently reported that systemic treatment with 25HC improved survival and reduced viral load, inflammation, and injury in the lungs of pigs during pneumonia with the coronavirus porcine reproductive and respiratory syndrome virus (8).

25HC is further oxidized by CYP7B1 on carbon-7, yielding the bioactive oxysterol 7α,25HC. 7α,25HC is reportedly depleted in *Ch25h*-null mice (24) and generated in response to exogenous 25HC substrate (2), and may regulate trafficking of immune cells to the lung in some settings (4, 6, 11, 12, 32). Given this, we tested mice null for the 7α,25HC receptor, EBI2 (encoded by *Gpr183*), in the SARS-CoV-2 pneumonia model in order to exclude possible confounding effects by this downstream lipid in our studies. We did not find any change in lung viral load in *Gpr183*^−/–^ mice nor alterations in body weight loss in EBI2 inhibitor-treated mice out to day 5 p.i., suggesting that, at least at these early time points, 7α,25HC does not play a significant role in anti-SARS-CoV-2 host defense. Importantly, mice deficient in EBI2 mount defective T-cell-dependent plasma cell and germinal center responses (24, 33, 34) and thus the small increase in viral titers on d5 may be related to defects in anti-viral T and B cell responses with EBI2 inhibition.

The increase in 25HC-treated animals of BALF albumin, an established metric of pulmonary microvascular lung injury (18), suggests a 25HC effect upon the pulmonary endothelial barrier, a putative cellular target of SARS-CoV-2 (22). 25HC is reportedly cytotoxic to endothelial cells (35-38), perhaps explaining our finding. The elevated plasma cytokines we observed in 25HC-treated mice may also derive from endothelial cells, given prior reports that 25HC augments cytokine expression by endothelial cells (39). Future studies will be necessary to examine the effect of 25HC on pulmonary vascular biology in greater detail. One group recently reported that direct intratracheal instillation of 25HC reduces LPS-induced lung inflammation (40). While this route might be expected to improve delivery to the alveolar epithelium and to reduce exposure of the endothelium as compared to i.p. injection, intratracheal aspiration is well-known to yield highly patchy, heterogenous deposition in the lung (41). Aerosol delivery reportedly provides superior alveolar delivery (41), but methods for aerosolization of 25HC have not been reported, to our knowledge.

Although our findings suggest that caution is warranted in extrapolating from studies of 25HC in other viruses to SARS-CoV-2, we propose that further exploration is still warranted of strategies that deliver 25HC more selectively to the primary cellular target of infection, respiratory epithelial cells. Along these lines, formulation of 25HC with cationic lipid as nanovesicles was recently reported to significantly improve lung-selective targeting of systemically dosed 25HC (42). Additional emerging strategies, such as conjugation of 25HC to viral fusion inhibitory peptides (13), also warrant future investigation in SARS-CoV-2 pneumonia. Given that CH25H and 25HC are reportedly increased in chronic obstructive pulmonary disease patients (4, 43) and in obese subjects (44), future studies to define the impact of native oxysterols on COVID-19 risk in lung disease (45) and in obesity (46) will also be of interest.

## Abbreviations

BALF: bronchoalveolar lavage fluid
Ch25h: cholesterol-25-hydroxylase
25HC: 25-hydroxycholesterol

## Data Availability

All data are contained within the manuscript.

## Acknowledgments

We are grateful to the staff of the NIAID ABSL2 and ABSL3 facilities and the NIAID SARS-CoV-2 Virology Core.

## Figure Legends

**Figure E1.**
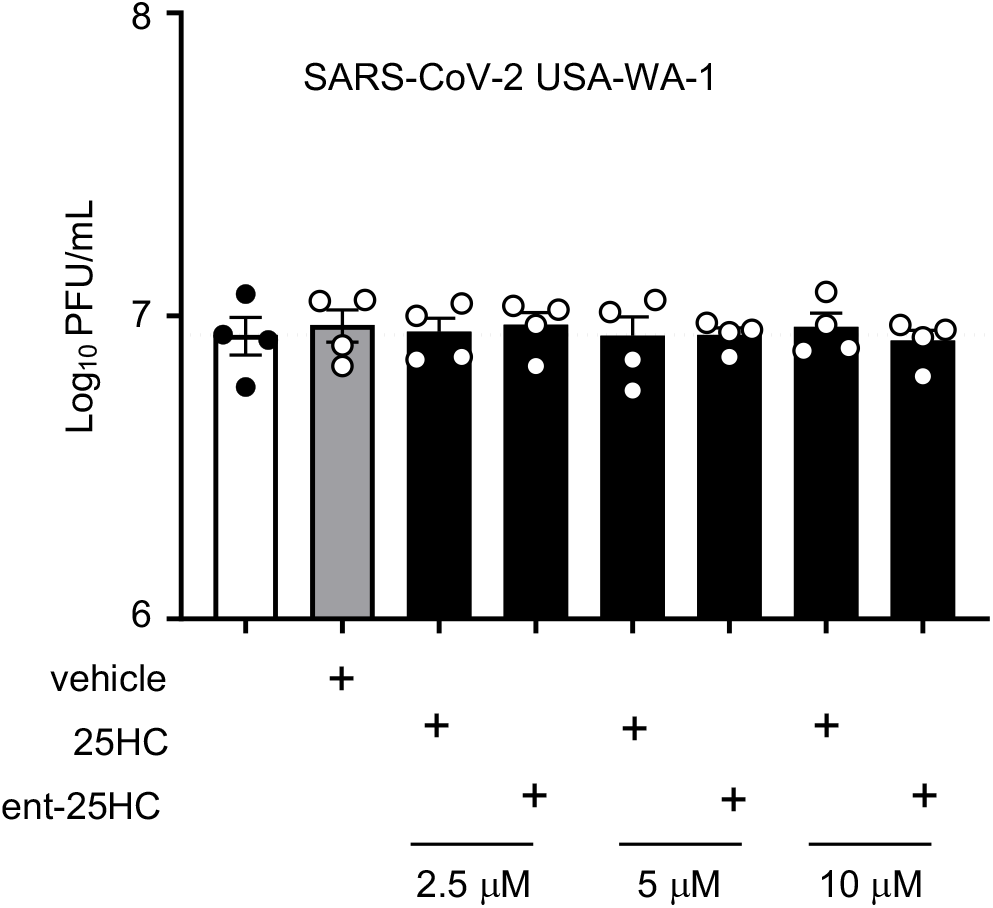
25-hydroxycholesterols do not antagonize SARS-CoV-2 as assessed by yield reduction assay. TMPRSS2-expressing VeroE6 cells were treated with vehicle or oxysterols for 24h as indicated, infected with SARS-CoV-2 USA-WA-1 for 1h, re-incubated in oxysterol, and then media harvested 24h later and evaluated by plaque assay. Data are mean +/- s.e.m. and are representative of 2-3 independent experiments.

**Figure E2.**
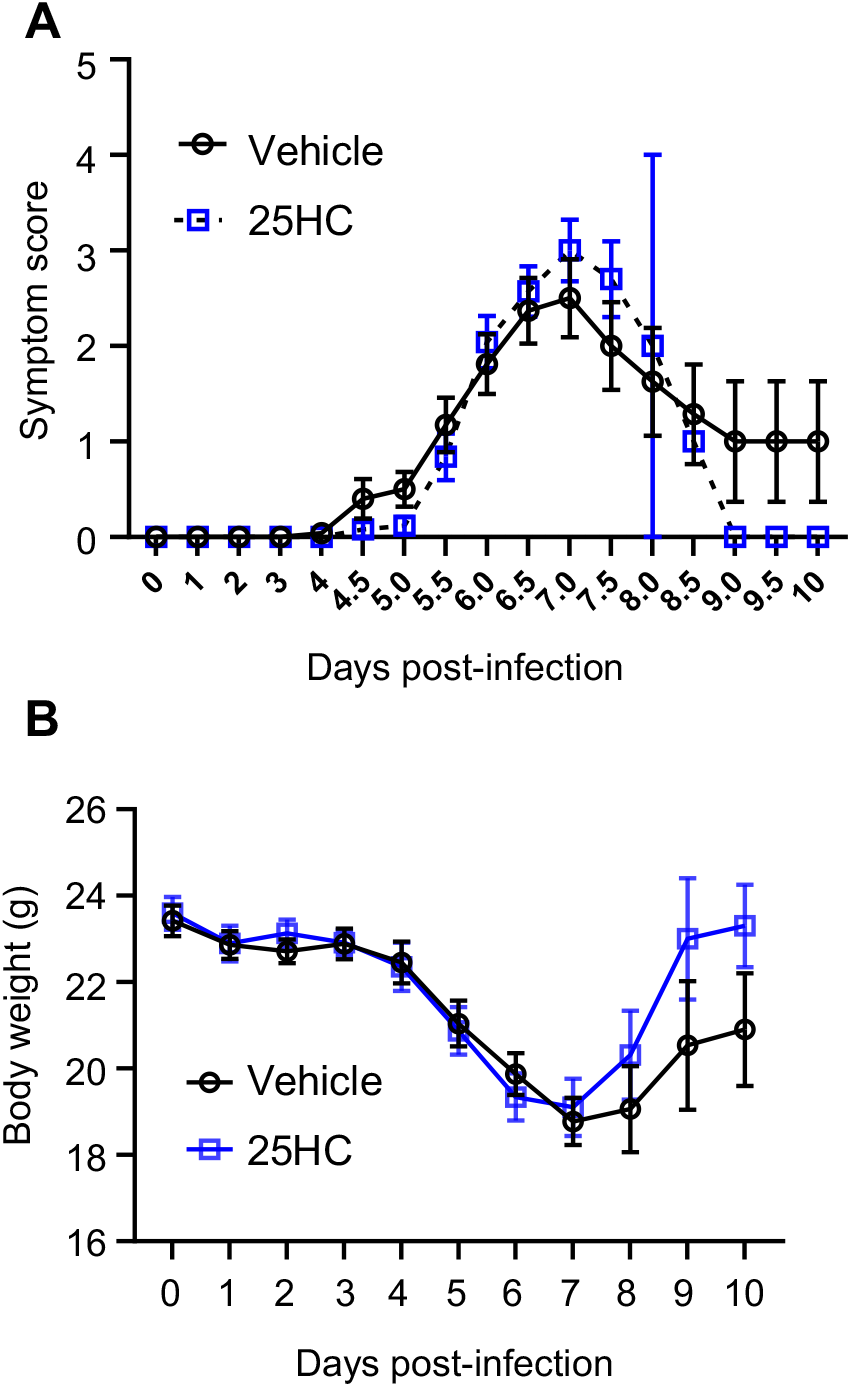
25HC treatment does not impact morbidity during murine SARS-CoV-2 pneumonia. K18-hACE2 mice were injected i.p. with 50 mg/kg/d 25HC (or vehicle) starting at -4h preceding i.n. inoculation with SARS-CoV-2. (**A**) Aggregate symptom score and (**B**) body weight was monitored over 10 days p.i. N=25/treatment. p=n.s. for analyses in both *A* and *B*.

